# On the Social Structure behind Biological Collections

**DOI:** 10.1101/341297

**Authors:** Pedro C. de Siracusa, Luiz M. R. Gadelha, Artur Ziviani

**Author notes:** {, }.

## Abstract

In this paper we describe two network models as a base for understanding the relevance of social processes involving collectors for shaping the composition of biological collections. **Species-Collector Networks** (SCNs) represent the interests of collectors towards particular species, while **Collector CoWorking Networks** (CWNs) represent collaborative ties between collectors during fieldwork. We demonstrate the practical use of our models with species occurrence data from the University of Brasília Herbarium.

## 1. Introduction

Biological collections are regarded as invaluable sources of primary biodiversity information, storing a large amount of physical materials and digital records that document the history of existence of organisms on geographic space. Data from biological collections have been increasingly used for a multitude of ecological and conservationist investigations, including the prediction of species responses to climate change, the selection of areas of high priority for conservation, the construction of red lists of threatened species, and the study of routes of biological invasion, just to cite a few [Nualart et al. 2017, Chapman 2005].

Many of the aforementioned studies involve the investigation of biogeographical patterns of species using **species occurrence records** (Figure 1), regarded as a fundamental source of biodiversity information. Each species occurrence record represents a *gathering act*, in which a *collector* (or team of collectors) collects a material evidence of the existence of a species at some locality and at some point in geographical space and time. Such an evidence is referred to as a *specimen* (not to be confused with *species*). After the specimen is properly incorporated to a collection, its taxonomic identity can be determined by professional taxonomists. This process consists of assigning a scientific name to the specimen, at the highest taxonomic rank as possible (usually species).

**Figure 1.**
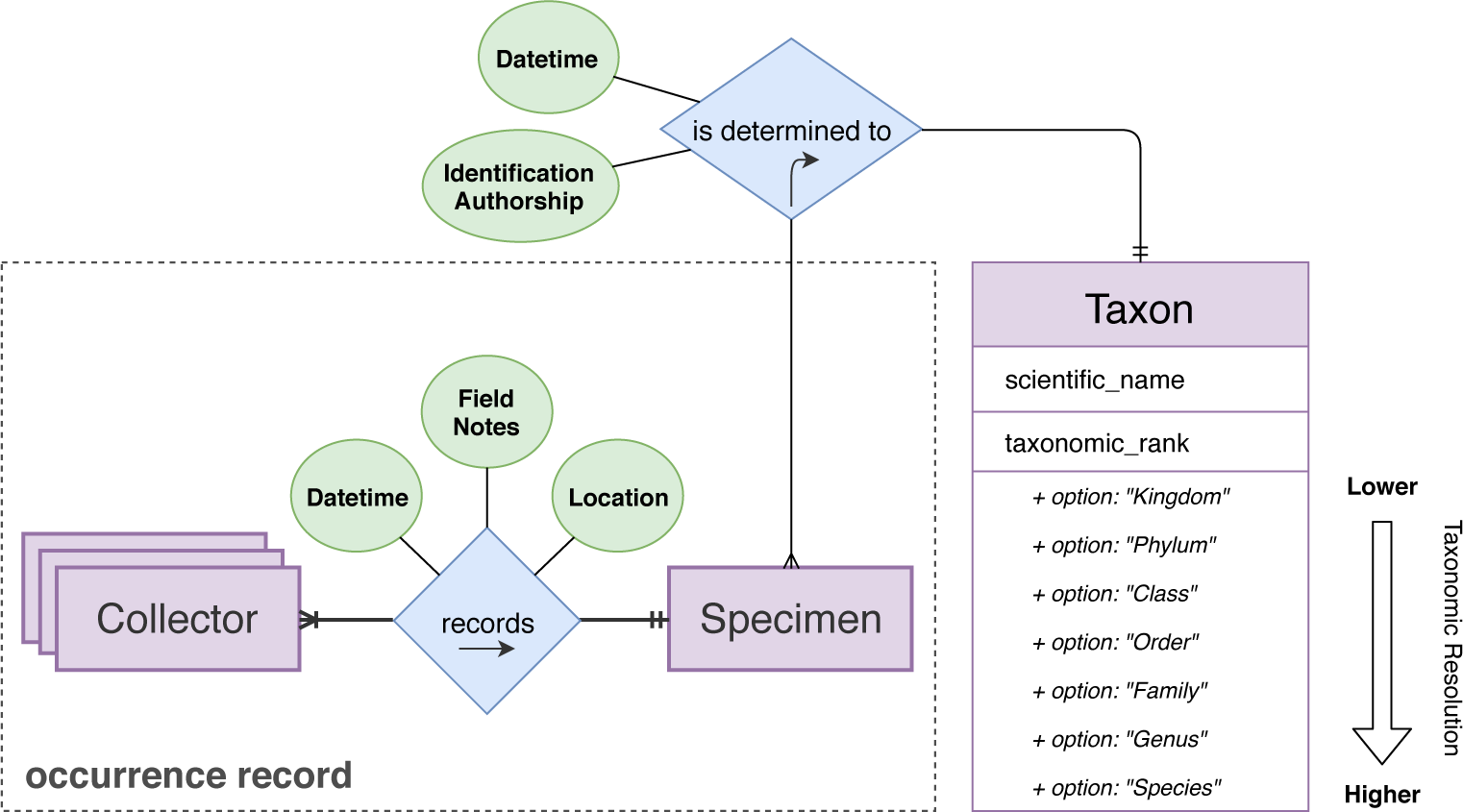
Entity-relationship diagram illustrating the main features of species occurrence records. The cardinality of relationships is represented using the Crow’s foot notation.

One important caveat of biological collections is that they are composed of occurrence records gathered at an enormous variety of circumstances, by a large community of collectors with distinct interests and using distinct sampling strategies. Collectors also tend to oversample species of their direct interests, at their preferred locations. As a consequence, biological collections fail to accurately represent the biological diversity within their actuation regions.Instead, they provide a partial sampled view, biased towards the interests and habits of their most productive collectors.

In this context, we propose a network-based approach for understanding how complex arrangements of the perceptions, interests and interactions of collectors shape the assemblage of biological collections. The approach is based on two classes of network models structuring collaborative relations involving pairs of collectors; and interest relations involving collectors and the species they record.

## 2. Network Models

Network models have been used in a myriad of contexts for the investigation of complex systems of interacting entities, from documents referring to each other in the World Wide Web [Barabási and Albert 1999] to organisms interacting in ecological webs [Bascompte 2007]. Networks are mathematically structured as *graphs*, composed of *nodes* (representing entities) and *edges* (representing links between pairs of entities). In this section we describe two classes of network models for investigating the taxonomic interests and collaborative behavior of collectors.

### 2.1. Species-Collector Networks

Species-Collector Networks (SCNs) are a particular type of *interest network*, representing the interests of *collectors* towards *species* they have recorded in field (Figure 2). Interest relationships necessarily involve a species and a collector, and can be semantically described as “**collector** samples **species**” or, conversely, “**species** is sampled by **collector**”. As the network is composed of two distinct classes of entities which necessarily interact with each other, the network is formally represented as an *undirected bipartite graph*

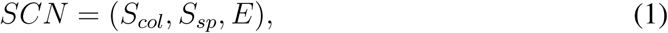

where *S_col_* and *S_sp_* are disjoint sets of nodes representing collectors and species, respectively; and E is the set of undirected edges connecting members of *S_col_* to *S_sp_*. The *bipartite constraint* ensures that all edges necessarily connect nodes from distinct sets, thus avoiding the introduction of semantic inconsistencies in the model.

**Figure 2.**
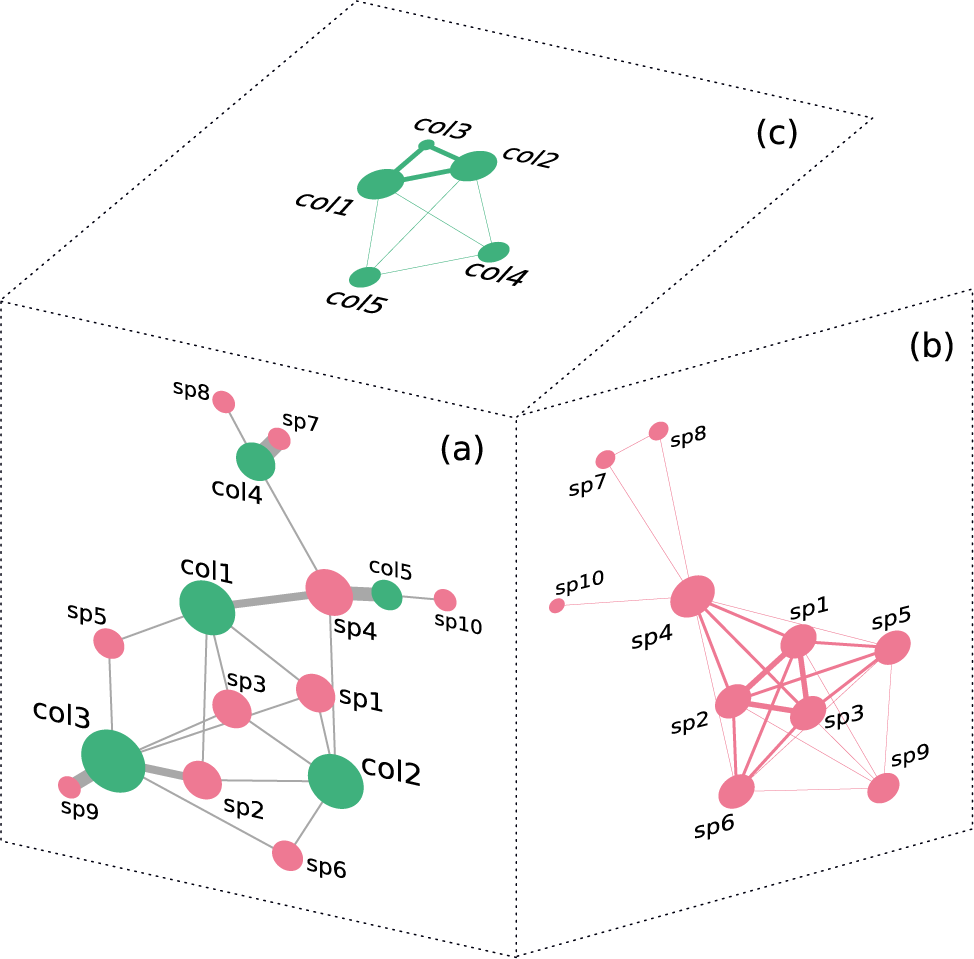
Multiple perspectives of a Species-Collector Network (SCN): (a) Unprojected bipartite network, showing collectors (green) linked to the species they have recorded (red); (b) Projection onto the species set; (c) Projection onto the collector set. In this figure, node size reflects the degrees of species and collectors, and edge thickness reflects the strength of the links.

The network is constructed from a species occurrence dataset, requiring information on (i) the names of the collectors and (ii) the species identity of the specimen at each record. Each collector in the team is linked to the recorded species, and the total number of times each collector or species occurs in the dataset is stored in their respective nodes as the *count* attribute. Edges are weighted proportionally to the number of times each species-collector association is observed in the dataset.

It is often convenient to summarize bipartite graphs into one-mode graphs, by linking together nodes from the same set if they are intermediated by a node from the opposite set. This operation is called a *bipartite projection*, and can be performed over each set of nodes of a bipartite graph. A projection over the *S_sp_* set provides a *species-centric* perspective (Figure 2(b)), in which species are directly connected if they have been recorded by at least one collector in common. A complementary *collector-centric* perspective (Figure 2(c)) is obtained by performing a projection over the *S_col_* set, such that collectors are directly connected if they have recorded at least one species in common. At each projection, the weight of edges connecting nodes from the same set is proportional to the similarity of their neighborhoods in the unprojected bipartite graph (i.e., the composition of nodes from the opposite set to which they are linked).

Another relevant operation defined for SCNs is the *taxonomic aggregation*. This process simplifies the network by grouping species nodes into taxa^1^ at higher taxonomic ranks (e.g. genus or family). Therefore, taxonomic aggregation reduces the number of nodes from the *S_sp_* set, with the side-effect of increasing network density^2^. The *taxonomic resolution* of a SCN is thus equivalent to the taxonomic rank at which species are aggregated in the model.

### 2.2. Collector CoWorking Networks

Collector CoWorking Networks (CWNs) are a particular type of *collaboration network*, structured from *collaboration* (or *coworking*) relationships between *collectors* who collect *specimens* together in field (Figure 3). Each coworking tie involves a pair of collectors, and can be semantically described as “**collector***_a_* records a specimen with **collector***_b_*”. As opposed to SCNs, the taxonomic identities of the specimens recorded are not represented in CWNs. The network is composed of nodes from a single entity class, being formally represented as an *undirected graph*
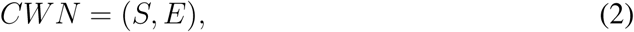

where *S* is the set of nodes representing collectors and *E* is the set of undirected edges connecting members of *S*.

**Figure 3.**
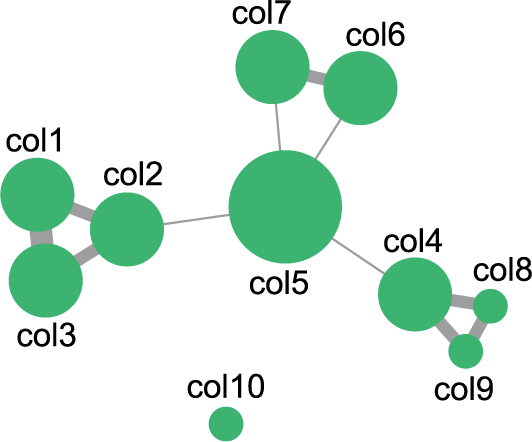
General aspect of a Collector CoWorking Network (CWN). Node size reflects the number of records a collector has authored, and edge thickness reflects the number of collaborative records coauthored by a pair of collectors.

This network is also constructed from species occurrence datasets, though it only requires information on the names of the collectors that have co-authored each record. For each record with team size *n*, 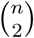 links connecting all collectors in a pairwise fashion are either formed or strengthened. Similarly to SCNs, the *count* attribute of each collector node stores the number of times the collector is observed in the dataset. Edge weight represents the strength of each link between collectors, and is calculated based on some user-defined weighting rule. We have obtained good results by adopting the *hyperbolic weighting rule* [Newman 2001], which assumes that collaborative ties derived from interactions in smaller teams tend to be stronger.

## 3. Proof of Concept: the University of Brasília Herbarium

In order to demonstrate the applicability of our models, we explored the social structure and interests of collectors from the University of Brasília Herbarium (UB) [Munhoz et al. 2018]. We used the species occurrence dataset of the UB herbarium, made available through the GBIF^3^ platform, for building both the SCN and CWN models. In order to improve the readability of the figures^4^, we aggregated the SCN at the family rank, and filtered out some less relevant nodes, which we considered irrelevant for the interpretation of the networks.

A first aspect to notice is that few collectors in the SCN have recorded a very high number of families (e.g. ‘*irwin,hs*’), while the majority of them have recorded only a few. Conversely, most families have been collected by few collectors, while some of them (e.g. *Fabaceae*) are collected by many. The central region of the SCN (Figure 4) contains some of the most relevant collectors of the herbarium (including ‘*irwin,hs*’ and ‘*heringer,ep*’), as well as very representative families of the Cerrado biome (*Fabaceae*, *Myrtaceae*, *Asteraceae*). Families at the central region are those more widely collected, while those towards the extremities are recorded by more specialized groups of collectors. We visually identified three main **communities of common interests**, formed by groups of collectors who are more specialized towards particular subsets of families than are other collectors, external to the group. Nodes within polygon *i* comprise a large part of the collectors from the *Cryptogams Lab*^5^ (headed by ‘*camara,peas*’, ‘*carvalhosilva,m*’ and ‘*souza,mgm*’), together with the families they are more interested in (especially *Sematophyllaceae*, a family of mosses). Polygon *ii* contains green algae collectors (‘*leite,alta*’, ‘*castelobranco,cw*’ and ‘*grando,jv*’, mostly recording family *Desmidiaceae*); and polygon *iii* contains another member of the *Cryptogams Lab* (‘*souza,mgm*’), who is particularly interested on diatoms, especially from families *Eunotiaceae*, *Naviculaceae* and *Pinnulariaceae*. However, collectors composing communities of common interests do not necessarily also collaborate during field collections. This aspect can be investigated from the CWN.

**Figure 4.**
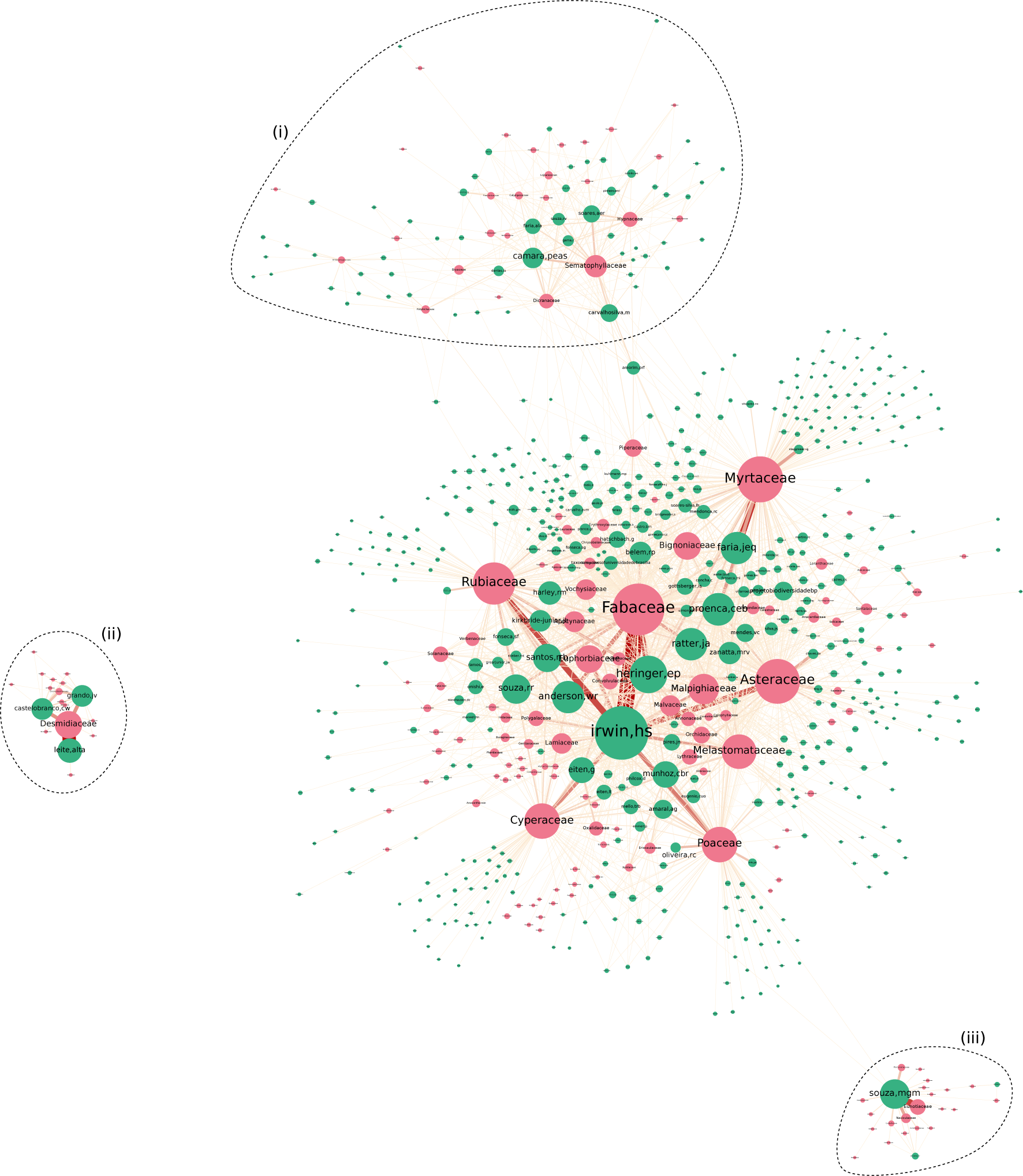
General aspect of the UB SCN, taxonomically aggregated at the family rank. Species and collector nodes are colored in pink and green, respectively. Node size is based on the *count* attribute, whilst the color and width of edges reflect their weight. Polygons (i), (ii) and (iii) are placed around communities that are visually most distinguishable.

Similarly to what we observed for the SCN, few collectors have collaborated with many others in the CWN (e.g. *‘proenca,ceb’*, at the center of the network), while the majority of them hold collaboration ties with quite a few colleagues (Figure 5). We identified a total of 30 **coworking groups** in the CWN, by using the *Louvain algorithm* for community detection [Blondel et al. 2008]. In some cases, collectors forming communities of common interests also form coworking groups (as it is the case of the *Cryptogams Lab* team, colored in pink, except for ‘*souza,mgm*’). It also happens that collectors belong to the same community of common interests but do not collaborate (e.g. ‘*bringel,jba*’ and ‘*chaves,da*’, specialists in family *Asteraceae*); or that collectors belong to the same coworking group but have very distinct taxonomic interests (e.g. ‘*faria,jeq*’ and ‘*zanatta,mrv*’). Finally, coworking groups can only be formed between collectors whose periods of collecting activities overlap. For instance, collectors from the green group (around ‘*irwin,hs*’) are considered to be the founders of the herbarium, with activities ranging from years 1960 to 1980, while a considerable part of collectors in network became active since 1990. Thus, coworking groups can also be formed on a temporal basis.

**Figure 5.**
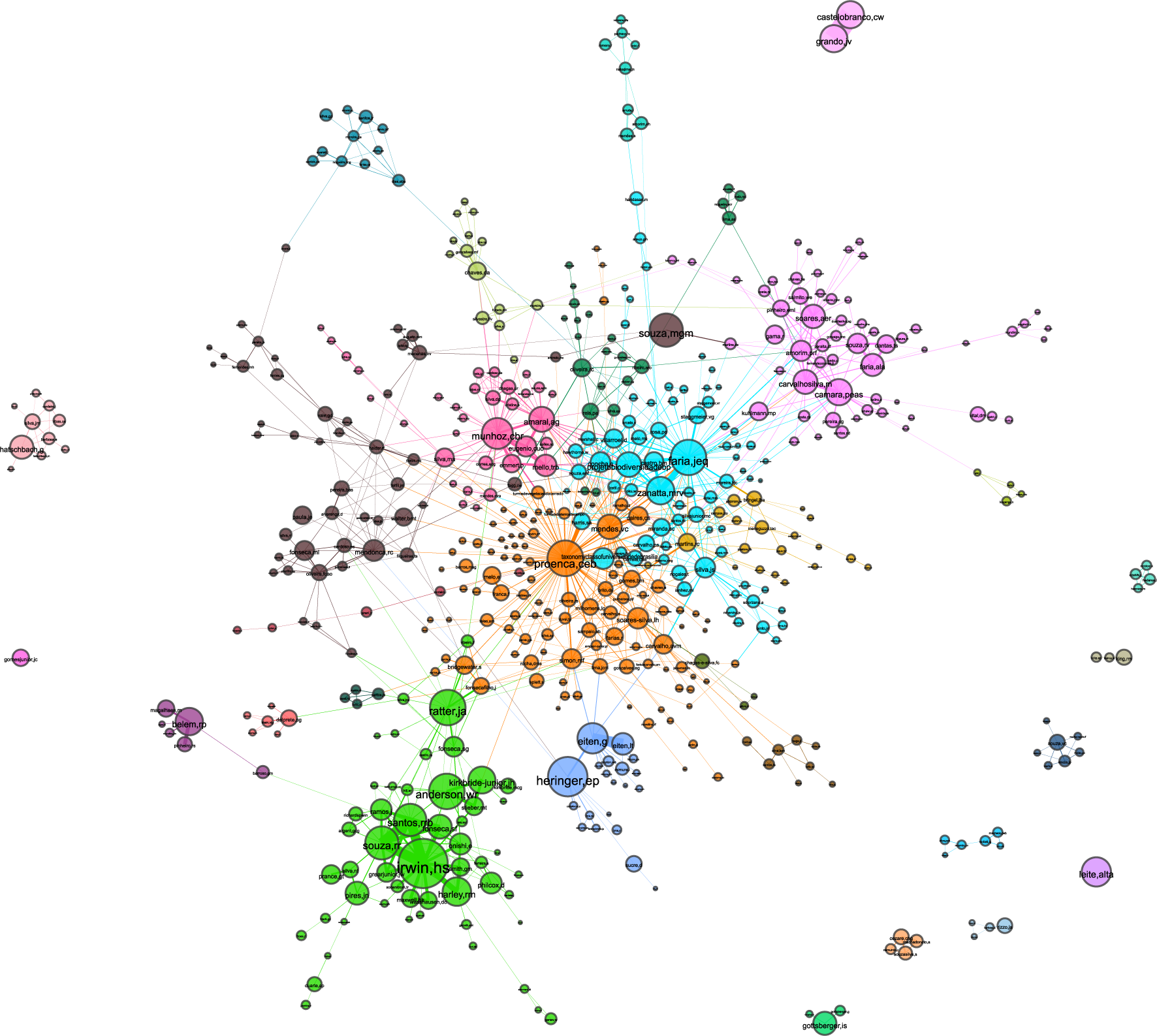
General aspect of the UB CWN. A total of 30 distinct coworking groups have been identified, and are differentiated by color. Node size is based on the *count attribute*, whilst the width of edges reflext their weight.

## 4. Final Remarks and Perspectives

Collaboration networks have been widely adopted in scientometric studies for investigating the patterns of collaboration within scientific communities [Newman 2004]. To the best of our knowledge, however, no previous attempts to investigate how collector social structures shape the composition of biological collections have been reported in scientific literature. The most similar study that we are aware investigates the formation of botanical exchange clubs from the 19*th* and 20*th* centuries in Britain and Ireland [Groom et al. 2014]. Such clubs were composed of botanists who corresponded to each other by exchanging plant specimens, with the goal of expanding their collections. Another recent study uses network analysis to investigate the connectivity patterns and roles of organizations in the Biodiversity Informatics landscape [Bingham et al. 2017]. Another interesting study, which is at some level analogous to our SCNs, uses a network-based approach for modeling the interests of music listeners towards genera, with the goal of characterizing collective music listening habits among users of media streaming platforms [Lambiotte and Ausloos 2006].

We believe our network models open new perspectives for research in the field of Biodiversity Informatics, especially for applications that rely on data from biological collections. With further developments from this work, we expect to provide mechanisms for systematically *profiling collectors*: classifying them according to their expertises, their collecting behaviors and their social roles in the collections they contribute to. Conversely, species can be also characterized based on the profiles of collectors who most record them. For instance, rare species tend to be mostly collected by experienced collectors, as they usually explore higher diversities of habitats while collecting [ter Steege et al. 2011]. We also believe these models provide the structural basis for a mechanism of *contextual enrichment of records*, allowing the investigation of how informative the composition of collector teams can be about the circumstances of the gathering act (e.g. an experienced collector teaming with many novices could indicate a recording taken during a field class).

## Acknowledgements

This work was partially supported by CAPES, CNPq, FAPERJ, and FAPESP. In particular, authors acknowledge the INCT in Data Sciences (CNPq no. 465560/2014-8). Authors are also grateful to Marinez Siqueira and Eduardo Dalcin, biodiversity researchers from the Research Institute Botanic Garden of Rio de Janeiro, Brazil, who provided insightful comments along the development of this work.

## Footnotes

1 Plural of taxon.

2 Network density is a measure of how close a network is to being fully connected.

3 https://www.gbif.org/

4 Interactive versions of the networks are available at https://lncc-netsci.github.io/pedrocs/networks/

5 http://labcriptounb.blogspot.br

